# Genome-Wide Profiling of Soybean WRINKLED1 Transcription Factor Binding Sites Provides Insight into Seed Storage Lipid Biosynthesis

**DOI:** 10.1101/2024.01.23.576967

**Authors:** Leonardo Jo, Julie M. Pelletier, Robert B. Goldberg, John J. Harada

**Author notes:** To whom correspondence should be addressed. Email: John J. Harada, Department of Plant Biology, 605 Hutchison Drive, University of California, Davis, CA 95616, USA; Leonardo Jo, Experimental and Computational Plant Development, Utrecht University, Padualaan 8, 3584 CH Utrecht, The Netherlands; and Robert B. Goldberg, Department of Molecular, Cell, and Developmental Biology, 4121 Terasaki Research Building, University of California, Los Angeles, CA 90095, (310) 825- 9093. These authors contributed equally to this work (Leonardo Jo, Julie M. Pelletier). Current address, Experimental and Computational Plant Development, Utrecht University, Padualaan 8, 3584 CH Utrecht, The Netherlands.

## Abstract

Understanding the regulatory mechanisms controlling storage lipid accumulation will inform strategies to enhance seed oil quality and quantity in crop plants. The WRINKLED1 transcription factor (WRI1 TF) is a central regulator of lipid biosynthesis. We characterized the genome-wide binding profile of soybean (Gm)WRI1 and show that the TF directly regulates genes encoding numerous enzymes and proteins in the fatty acid and triacylglycerol biosynthetic pathways. GmWRI1 binds primarily to regions downstream of target gene transcription start sites. We showed that GmWRI1 bound regions are enriched for the canonical WRI1 DNA binding element, the AW Box (CNTNGNNNNNNNCG), and another DNA motif, the CNC Box (CNCCNCC). Functional assays showed that both DNA elements mediate transcriptional activation by GmWRI1. We also show that GmWRI1 works in concert with other TFs to establish a regulatory state that promotes fatty acid and triacylglycerol biosynthesis. In particular, comparison of genes targeted directly by GmWRI1 and by GmLEC1, a central regulator of the maturation phase of seed development, reveals that the two TFs act in a positive feedback subcircuit to control fatty acid and triacylglycerol biosynthesis. Together, our results provide new insights into the genetic circuitry in which GmWRI1 participates to regulate storage lipid accumulation during seed development.

**Significance Statement:** We report the genome-wide profiling of DNA sequences bound by and the genes directly-regulated by soybean WRINKLED1, a central regulator of storage lipid accumulation in oilseed plants. The information offers new insights into the mechanisms by which WRINKLED1 regulates genes encoding lipid biosynthetic enzymes and establishes a regulatory environment that promotes oil accumulation, and it may aid in the design of strategy to alter storage lipid accumulation in oilseeds.

## Introduction

Oils derived from plants are used primarily for food-based applications, although they also have broad utility in industrial applications (1). Given that lipids are essential for human nutrition and that biofuels are of increasing importance as a sustainable energy source, the demand for plant oils is anticipated to increase. Because oilseeds account for approximately 55% of world vegetable oil production (2), developing crops with increased and/or modified seed oil content may be instrumental in meeting these demands. To achieve this goal, it is crucial that we understand the regulatory mechanisms that underlie seed storage lipid accumulation to inform the development of biotechnological tools that will improve the quality and quantity of seed oils.

Fatty acids (FAs) are stored primarily in the form of triacylglycerol (TAG) in seeds. As summarized in Fig. 1A, the FA biosynthetic pathway in seeds occurs in plastids (3, 4). The first committed and rate-limiting step in FA biosynthesis is catalyzed by the heterotetrameric acetyl-CoA carboxylase (ACCase) enzyme that produces malonyl-CoA. Subsequent steps are catalyzed by fatty acid synthase, a multisubunit complex consisting of 10 distinct enzymes activities, that catalyzes the transfer of malonyl-CoA to acyl carrier protein (ACP), the sequential elongation of the acyl-ACP chain, and the release of non-esterified C_16_ or C_18_ fatty acids. In addition to the FA biosynthetic enzymes, several other enzymes responsible for the elongation and desaturation of FAs account for the diversity of FAs found in the seeds (5).

**Figure 1.**
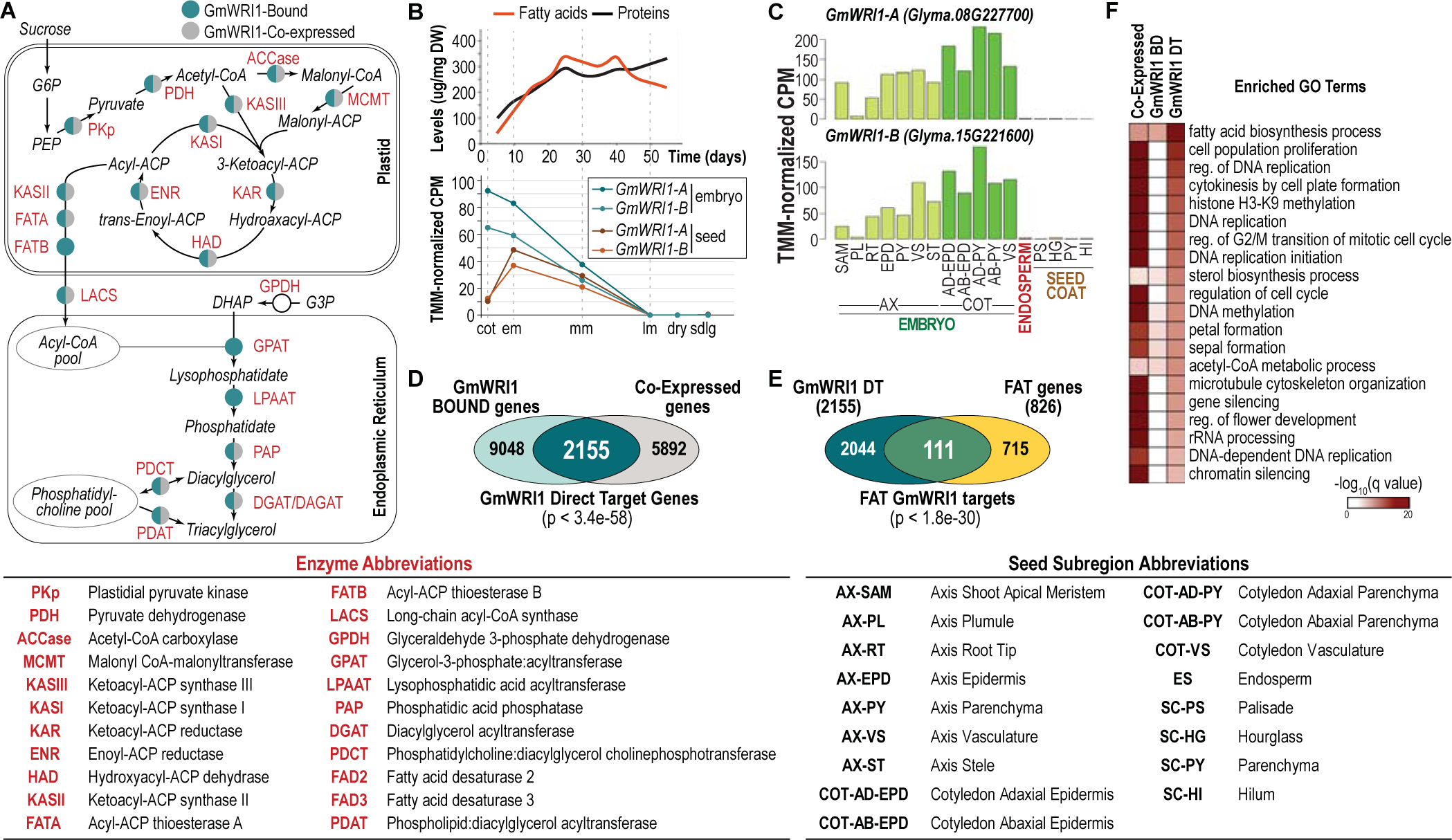
GmWRI1 acts as a central regulator of fatty acid and triacylglycerol biosynthesis in soybean seeds. (A) Simplified schematic diagram of FA and TAG biosynthesis from Baud and Lepiniec (11). Enzymes (labeled in red) encoded by genes that are bound by (teal) and/or co-expressed with (grey) GmWRI1. (B) Temporal changes in relative storage fatty acid and protein levels (top, adapted from Collakova *et al*. (56)) and of *GmWRI1-A* (*Glyma.08G227700*) and *GmWRI1-B* (*Glyma.15G221600*) mRNA levels (bottom, from the Harada Embryo mRNA- Seq Dataset, GEO Series GSE99571) during soybean embryo development. (C) mRNA levels of *GmWRI1-A* and *GmWRI1-B* in seed subregions of the embryo axis, and cotyledons, endosperm, and seed coat at the early maturation stage (GEO Series GSE46096). (D) GmWRI1 DT genes in soybean embryos at the early maturation stage. Venn diagram shows the overlap of GmWRI1 bound and coexpressed genes. Statistical significance (hypergeometric distribution) of the overlap between bound genes and co-expressed genes is indicated. (E) Venn diagram showing the overlap of the GmWRI1 DT genes at the em stage and of FAT genes, with the statistical significance of the overlap indicated (hypergeometric distribution). (F) Heatmap showing the q value significance of GO term enrichment for GmWRI1 co-expressed, bound, and GmWRI1 DT genes. The GO terms listed are the top 20 most enriched biological process GO terms for GmWRI1 DT genes. Abbreviations: cotyledon (cot), early maturation (em), mid-maturation (mm), late maturation (lm), seedling (sdlg).

TAG synthesis occurs in the endoplasmic reticulum through one of two pathways (6, 7). The acyl CoA-dependent pathway, known as the Kennedy pathway, catalyzes the sequential addition of fatty acid moieties to a glycerol-3-phosphate backbone to generate TAGs. In the alternative pathway, the acyl-CoA-independent pathway which predominates in soybean, fatty acids are first incorporated into the membrane lipid phosphatidylcholine prior to their incorporation into TAG. TAGs are stored largely in oil bodies that bud off from the endoplasmic reticulum and are surrounded by a phospholipid monolayer (8). Storage lipid and protein accumulation occurs primarily during the maturation phase of seed development (Fig. 1B) which follows the morphogenesis phase during which embryonic domains and tissues are established (9).

Thus, storage lipid accumulation in seeds requires the coordinated activity of many diverse enzymes that operate in distinct cellular compartments. Oilseed crops are particularly efficient in FA and TAG s biosynthesis, with storage lipids accounting for more than 60% of seed dry mass (10). This massive accumulation of TAGs and the observation that FA biosynthesis genes show similar patterns of expression that peaks during the onset of the maturation program and declines during the later stages of maturation suggests that FA and TAG biosynthesis is precisely regulated during seed development (11). Several reports indicate that FA biosynthesis is regulated primarily at the transcriptional level (12–15).

The major regulator of seed storage lipid biosynthesis is the WRINKLED1 (WRI1) transcription factor (TF), a member of the APETALA2 TF family (16). WRI1 has been shown to be both necessary and sufficient for FA accumulation in many oilseed crops. For example, Arabidopsis (*at*)*wri1* loss-of-function mutants exhibit an 80% reduction in seed oil content (17), and ectopic expression of *WRI1* results in significant increases of FA-related gene expression and, consequently, increased oil content in several plant species (16, 18–22).

WRI1 affects storage lipid accumulation by regulating the expression of genes involved in FA biosynthesis. Experimental modulations of *WRI1* expression that alters seed oil levels cause corresponding changes in the expression of FA biosynthesis genes, including those encoding ACP, 3-ketoacyl-ACP synthases (KAS), and subunits of ACCase, suggesting that these genes are activated transcriptionally by WRI1 (19, 23, 24). Consistent with this hypothesis, the AW Box (CNTNGNNNNNNNCG), a DNA element bound by WRI1 *in vitro* (23), is enriched in FA biosynthesis genes.

Despite its key regulatory role, a detailed, mechanistic understanding of how WRI1 regulates storage lipid accumulation in seeds is lacking. In particular, a comprehensive, genome-wide description of all genes regulated by WRI1 has not been reported, limiting our understanding of the exact mechanism by which WRI1 acts to regulate seed oil accumulation.

To address this limitation, we analyzed the genome-wide binding profile of soybean WRI1 (GmWRI1) during seed development and provide the first comprehensive description of WRI1 binding sites and the genes directly regulated by WRI1 in plants. Our results provide new insights into the diverse mechanisms by which WRI1 acts as a central regulator of storage lipid accumulation in seeds.

## Results

### GmWRI1 Binds and Regulates Genes Involved in Fatty Acid Biosynthesis

We identified genes that are directly regulated by GmWRI1-A (Glyma.08G227700) and GmWRI1-B (Glyma.15G221600). Although there are several *WRI1* homologs in soybean, *GmWRI1-A* and *GmWRI1-B* showed the highest degree of DNA sequence similarity with *AtWRI1,* and other have demonstrated their ability to rescue the *atwri1* seed oil mutant phenotype in Arabidopsis (25). As summarized in Fig. 1B, data from existing RNA-Sequence (RNA-Seq) databases (GEO accession GSE99571) showed that both mRNAs are present at high levels in embryos at the cotyledon (cot) and early maturation (em) stages and decline at later stages of embryo development, preceding the onset of FA accumulation in soybean seeds. Analysis of the spatial distribution of *GmWRI1* mRNAs in em stage seeds (GEO accession GSE46096) showed that both accumulated predominantly in embryo subregions (Fig. 1C). The temporal and spatial expression patterns of *GmWRI1-A* and *GmWRI1-B* are consistent with their roles in controlling the FA and TAG biosynthetic programs in embryos.

To identify genes that are directly regulated by GmWRI1 during seed development, we first defined genes that were bound by the TFs. We performed chromatin immunoprecipitation followed by DNA sequencing (ChIP-Seq) experiments using em stage soybean embryos with antibodies that recognize both GmWRI1-A and GmWRI1-B (see Experimental Procedures).

GmWRI1 bound genes were defined as those with reproducible GmWRI1 ChIP-Seq peaks in a genomic region between 1 kb upstream of the gene transcription start site (TSS) to the transcription termination site. We identified 11,203 bound genes (Dataset S1). We also performed GmWRI1 ChIP-Seq experiments with cotyledon (cot)-stage embryos, because *GmWRI1* mRNA accumulation is high at that stage (Fig. 1B). However, the cot-stage results did not pass ENCODE quality thresholds for ChIP-Seq data unlike experiments at the em stage (26)(Dataset S2), as indicated by the smaller number of bound genes that were detected at the cot stage compared with the em stage (SI Appendix Fig. S1A, Dataset S1). Nevertheless, virtually all cot-stage GmWRI1 bound genes were also bound at the em stage (SI Appendix Fig. S1A).

Because most TF binding interactions do not represent transcriptional regulatory events (27), we filtered GmWRI1 bound genes for those that are co-expressed with the TF. The rationale is that a TF and the genes that it directly regulates are expressed in the same cell.

Given the *GmWRI1* mRNA spatial accumulation pattern, we defined co-expressed genes as those whose mRNA levels were at least two-fold or higher in embryo cotyledon parenchymal tissues compared to seed coat parenchyma and seed coat hilum tissues (FDR < 0.05, Harada–Goldberg Soybean Seed Development LCM RNA-Seq Dataset, GEO accession GSE99109; Dataset S1).

As shown in Fig. 1D, we identified 2,155 genes that were both bound by GmWRI1 and co-expressed with *GmWRI1*, a significant overlap (*P* < 3.4e-58), and we designated these as GmWRI1 direct target (DT) genes.

We compared GmWRI1 DT genes with Arabidopsis genes regulated by AtWRI1 to confirm that the soybean genes were WRI1 regulated. Because neither genes bound by AtWRI1 nor genes mis-regulated by *atwri1* mutations have been reported genome-wide, we identified genes that were differentially expressed (DE) in *atwri1-1* mutant embryos at the linear-cotyledon stage (Fig. 2A and S2). We found that 1,213 DE genes were downregulated (two-fold, FDR < 0.05) in *atwri1-1* compared with wild-type (WT) embryos, 848 of which correspond to annotated soybean genes (SI Appendix Fig. S3, Dataset S3). A significant number of these genes, 165 (P < 5.8e-14), overlapped with Arabidopsis homologs of GmWRI1 DT genes (1,690, Fig. 2C). To define Arabidopsis genes upregulated by ectopic *GmWRI1* expression, we transfected Arabidopsis leaf protoplasts with either the *GmWRI1-A* gene driven by the *CaMV35S* promoter (*p35S:GmWRI1*) or the *p35S:mCHERRY* gene (Fig. 2B). We identified 1,490 DE genes that were upregulated (two-fold, FDR < 0.05) in *35S:GmWRI1* versus *35S:MCHERRY* cells, 1097 of which correspond to annotated soybean genes (SI Appendix Fig. S3, Dataset S3). We found that 212 of these genes overlapped significantly with Arabidopsis homologs of GmWRI1 DT genes (P < 2.0e-17, Fig. 2D). Together, these results suggest that GmWRI1 regulates GmWRI1 DT genes. We designated the 401 genes (p < 1.7e-214) that are downregulated in *atwri1* mutant embryos and upregulated by *35S:GmWRI1* overexpression in leaf cells as AtWRI1 directly regulated (AtWRI DR) genes (Fig. 2E).

**Figure 2.**
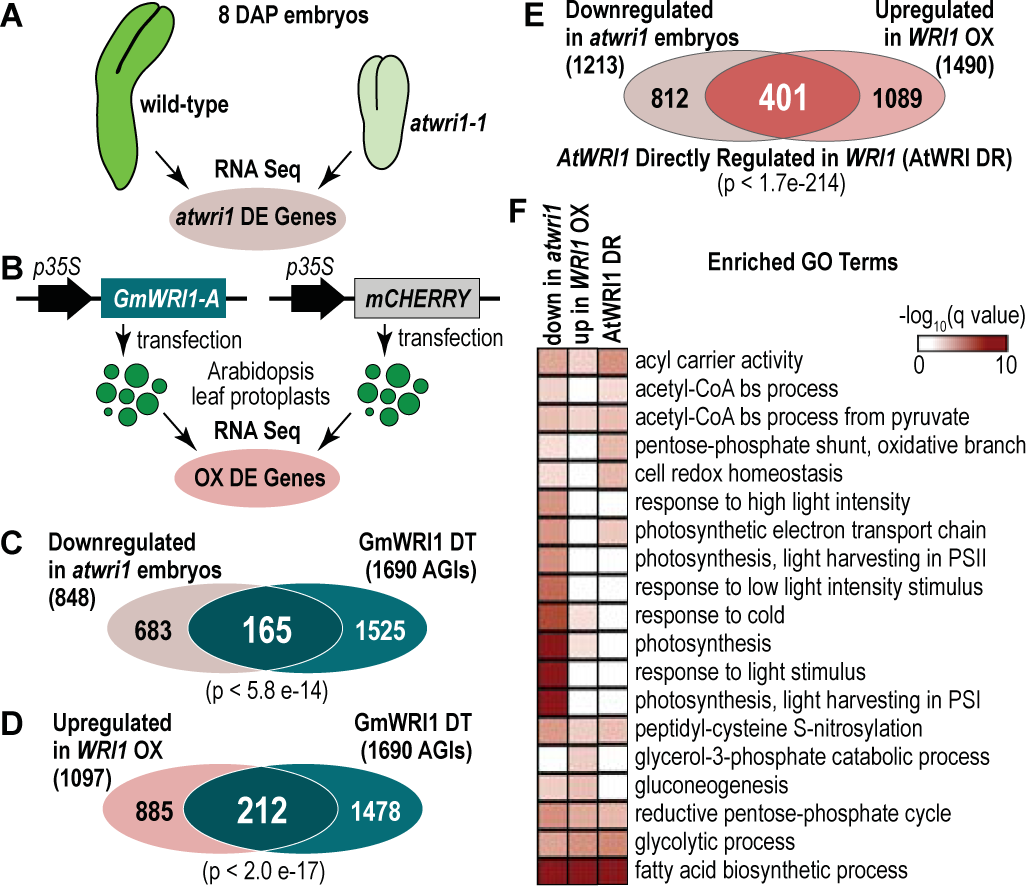
Identification of WRI1 directly regulated genes in Arabidopsis. (A) Overview of strategy used to identify Arabidopsis genes differentially expressed (A) in *atwri1-1* mutants compared with WT embryos at 8 days after pollination and (B) in leaf protoplasts transfected with *35S:GmWRI1* or *35S:mCHERRY* as a negative control. RNA-Seq experiments identified differentially expressed mRNAs that differed at least two-fold (FDR < 0.05) when compared to the control. (C) Venn diagram showing the overlap between genes downregulated in *atwri1* mutant embryos and GmWRI1 DT genes. (D) Venn diagram showing the overlap between genes upregulated in leaf protoplasts overexpressing GmWRI1 and GmWRI1 DT genes. (E) Venn diagram showing the overlap between genes downregulated in *atwri1* mutant embryos and upregulated in leaf protoplasts overexpressing *GmWRI1* to identify the AtWRI1 DR genes. (F) Heatmap showing the q value significance of enriched GO terms for downregulated genes in *atwri1* mutant embryos, upregulated genes in the *35S:GmWRI1* leaf cells, and of the AtWRI1 DR genes. GO terms listed are the top 10 most enriched biological process terms (q value < 0.01).

Two lines of evidence suggest that GmWRI1 DT genes are involved in FA and TAG biosynthesis. First, GmWRI1 DT genes were overrepresented for Gene Ontology (GO) terms involved with FA biosynthesis, such as fatty acid biosynthetic process and acetyl-CoA metabolic process, acetyl-CoA carboxylase activity, and fatty acid synthase complex (Fig. 1F, Dataset S1). Similarly, AtWRI DR genes were enriched for GO terms related to FA biosynthetic processes, such as fatty acid biosynthetic process, acyl carrier activity, and acetyl-CoA carboxylase activity (Fig. 2F, Dataset S3). GmWRI1 DT genes were also enriched for GO terms related to cell proliferation, such as cell population proliferation and cytokinesis by cell plate formation, opening the possibility that GmWRI1 may also regulate genes involved in processes other than storage lipid accumulation. Second, Fig. 1A shows that the vast majority of enzymes involved in FA and TAG biosynthesis are encoded by GmWRI1 DT genes, such as the ACCase subunits, biotin carboxylase (CAC2) and biotin carboxyl carrier protein (BCCP2), and KASIII, and in TAG assembly in the endoplasmic reticulum, such as phosphatidic acid phosphohydrolase (PAH), diacylglycerol acyltransferase (DGAT), and phosphatidylcholine diacylglycerol cholinephosphotransferase (PDCT). We note that two genes are bound by but not coexpressed with GmWRI1. To determine the extent to which GmWRI1 regulates genes involved in FA and TAG biosynthesis, we manually curated a list of FA and TAG (FAT) biosynthesis genes based on the Plant Metabolic Network (PMN) (https://plantcyc.org/databases/soycyc/8.0) and used the Arabidopsis Acyl-Lipid Metabolism (Aralip) (http://aralip.plantbiology.msu.edu/) to annotate these genes (Dataset S4). Of the 826 soybean FAT genes that encode different steps in the FA and TAG biosynthetic pathways, 111 were GmWRI1 DT genes (*P* < 1.8 e-30, Fig. 1E).

Together, our results confirm that GmWRI1 directly regulates genes required for storage lipid biosynthesis.

### WRI1 Primarily Binds Intragenic Regions Enriched for Specific DNA Motifs

To gain insight into the mechanisms by which GmWRI1 regulates FA and TAG biosynthetic genes transcriptionally, we analyzed bound regions in GmWRI1 DT genes. Surprisingly, we showed that the ChIP-Seq peaks which presumably represent GmWRI1 binding sites in cot and em stage embryos were mostly located downstream of the TSS in intragenic regions (Fig. 3, A and B, and S1b). This binding site profile differed from those of GmLEC1, GmAREB3, GmbZIP67, and GmABI3 TFs in soybean embryos whose ChIP-Seq peaks were mostly located in region upstream of the TSS (28, 29).

**Figure 3.**
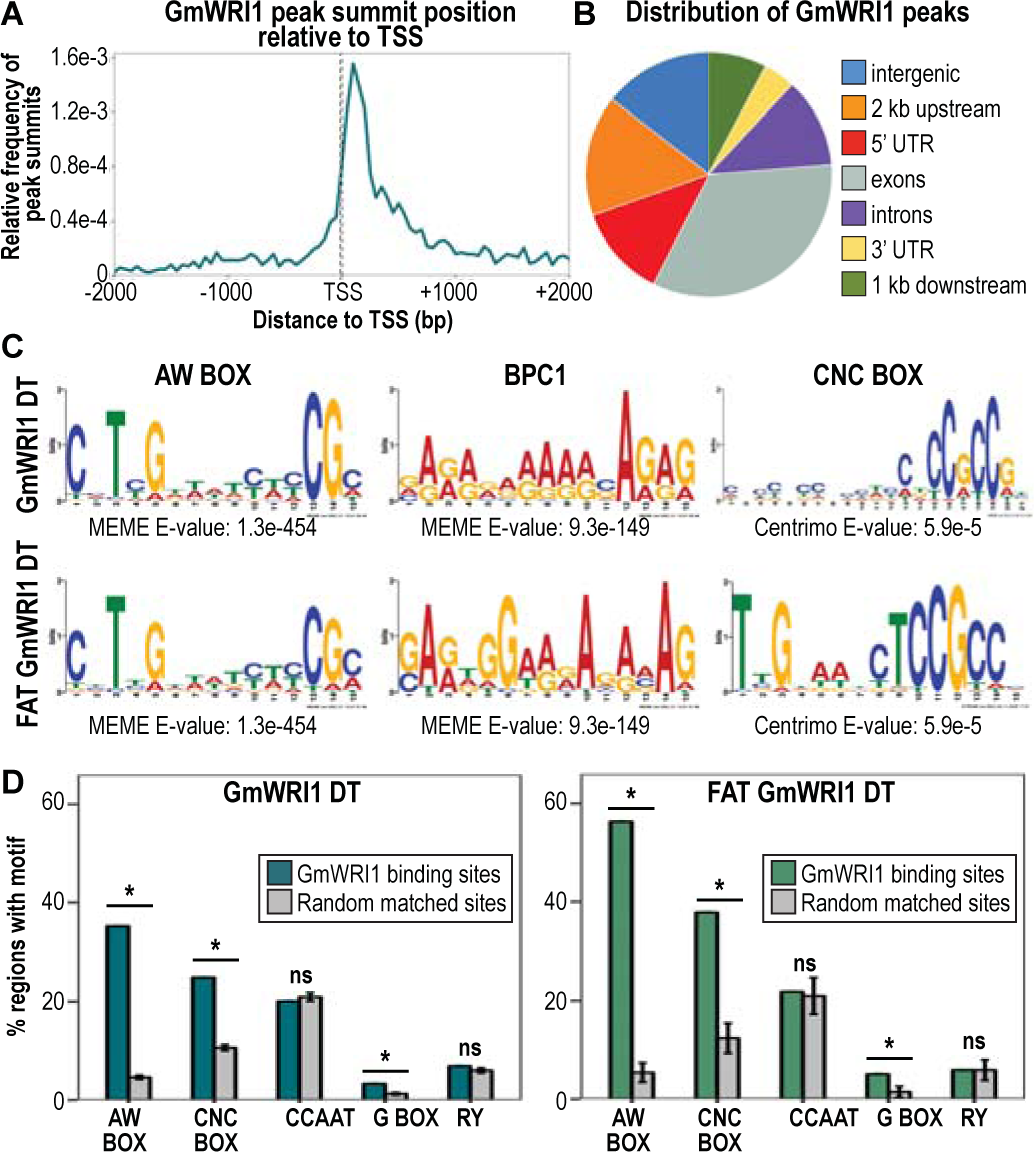
Binding profile and enriched DNA motifs in GmWRI1 binding sites. (A) Position of GmWRI1 ChIP-seq peak in reference to the TSS of GmWRI1 DT genes. (B) Pie chart showing the fraction of GmWRI1 ChIP-Seq peaks that overlap distinct genomic features (intergenic, 2 kb upstream of the TSS, 5’ UTR, exons, introns, 3’ UTR and 1 kb downstream of the TTS). (C) Position weight matrices of DNA motifs discovered *de novo* in GmWRI1 binding sites of GmWRI1 DT (top), and FAT GmWRI1 DT (bottom) genes, and their significance as indicated by their associated E values. (D) DNA motif enrichment in the binding sites of GmWRI1 DT (left) and FAT GmWRI1 DT (right) genes. Graphs show the percentage of bound regions (green) and randomly selected, equivalently sized and spaced regions (gray) containing the indicated DNA motifs. A Bonferroni-adjusted *P* value threshold of 0.01 was used to indicate the statistical significance (*) of enrichment, while n.s. indicates no significant difference.

#### AW Box DNA motif is highly enriched in GmWRI1 binding sites

We asked if DNA motifs were enriched in a 100 bp region surrounding the ChIP-Seq peaks, designated as GmWRI1 binding sites. One motif identified using *de novo* discovery analysis of GmWRI1 DT binding sites was the AW box motif, a DNA sequence previously shown to be bound by AtWRI1 *in vitro* (23)(Fig. 3C, Dataset S5). We showed that GmWRI1 DT and FAT GmWRI1 DT genes were significantly enriched for the AW Box (Bonferroni-adjusted *P* values < 0.01), with 35.2% and 56.3% of binding sites, respectively, containing full AW Boxes (Fig. 3D). These results suggest that the AW Box underlies the direct interaction of GmWRI1 with many, but not all, of its binding sites.

Because most GmWRI1 binding sites were located downstream of the TSS, we determined AW Box distribution in GmWRI1 DT and FAT GmWRI1 DT genes. AW Box motifs were found predominantly in regions downstream of the TSS, consistent with GmWRI1 binding site locations (SI Appendix Fig. S4, A and B). We note, however, that AW Box enrichment may be influenced by the higher GC content of regions downstream versus upstream of the TSS. Nevertheless, AW Box distribution and enrichment in GmWRI1 binding sites is consistent with a role for this DNA motif in specifying GmWRI1 occupancy in downstream regions.

#### Additional DNA motifs are enriched in WRI1 binding sites

*De novo* discovery analyses identified another DNA motif, CNCCNCC (designated CNC Box), that was enriched in the binding sites of GmWRI1 DT and FAT GmWRI1 DT genes (Fig. 3, C and D, Dataset S5). Although the CNC Box has not been shown to be bound by WRI1, its enrichment in GmWRI1 binding sites suggests its potential role in mediating GmWRI1 DT gene transcription. Consistent with this hypothesis, a similar motif was previously shown to be enriched in the 5’ UTR of FA-related genes in numerous plant species (30).

A GAGA-rich DNA motif was also discovered *de novo* in GmWRI1 binding sites (Fig. 3, B and C). This motif is similar to the BPC1 motif that is bound by the BASIC PENTACYSTEINE6 (BPC6) transcription factor which is proposed to recruit the Polycomb Repressive Complex (31). The BPC1 motif is found in the binding sites of many TFs that regulate the maturation phase of seed development, such as GmLEC1, GmAREB3, GmbZIP67 and GmABI3 (28, 29). Together, these results suggest that transcriptional regulation of GmWRI1 DT genes may involve DNA elements other than the well characterized AW Box.

Because FAs and TAGs are synthesized during the maturation phase, we asked whether the CCAAT Box (CCAAT), G Box (CACGTG) and RY (CATGCA) DNA elements that are known to be bound by the major maturation TF regulators, LEC1, bZIP67, ABI3, FUS3, and LEC2, were enriched in GmWRI1 binding sites (28, 32). These motifs were not identified by *de novo* discovery analysis, and only the G Box motif was significantly enriched in 3.3% and 5.0% of GmWRI1 DT binding sites and FAT GmWRI1 DT binding sites (Fig. 3D and Dataset S5).

This result suggests that GmWRI1-mediated transcriptional control of FA biosynthesis involves distinct cis-regulatory modules from those utilized by these other maturation TFs.

#### AW and CNC Box motifs are enriched in intragenic regions of AtWRI Directly-Regulated genes

We analyzed motif enrichment in AtWRI DR genes to determine if the motifs identified in GmWRI1 binding sites are conserved in Arabidopsis. AW Boxes and CNC Boxes were detected downstream of the TSS in 70.3% and 59.6%, respectively, of AtWRI DR genes (Fig. S5, Dataset S3). By contrast, the AW Boxes and CNC Boxes were detected in only 41% and 48.9%, respectively, of the downstream regions of 1000 sets of randomly matched genes (Dataset S3). In regions upstream of the TSS, the AW Box was enriched in 33.2% of AtWRI1 DR genes as compared with 18.5% in background regions (Fig. S5, Dataset S3), but no significant enrichment of the CNC Box was observed. These results are consistent with the observation that the AW Box and CNC Box are overrepresented in binding sites downstream of the TSS of genes regulated by WRI1, suggesting that these elements contribute to the transcriptional regulation of these genes.

### Functional Analysis of AW Box and CNC Box Motifs in GmWRI1 Direct Target Gene Binding Regions

We conducted transient assays to determine whether enriched AW Boxes and CNC Boxes are required for GmWRI1-mediated transcription of the four FAT genes shown in Fig. 4A: *ACP4* (Glyma.15G098500), *BCCP2* (Glyma.13G0 57400), *CAC2* (Glyma.05G221100), and *ACP3* (Glyma.19G240100). We cloned WT versions of the genes’ binding sites and mutant variants in which all AW Box (mAW), CNC Box (mCCNCC) or both (mAWmCNC) motifs were mutated (Fig. 4B, Dataset S6). Binding sites for the *ACP4*, *BCCP2,* and *CAC2* genes contained at least one full AW Box motif, whereas *ACP3* contained an abbreviated version of the AW Box (CNTNGNNNNNNCG) which was shown by others to be bound by AtWRI1 *in vitro* (33). With the exception of *BCCP2,* the binding sites all contained at least one CNC Box. Binding sites were fused upstream of a firefly *Luciferase* gene with a *35S* minimal promoter in a plasmid that also contained a *35S:Renilla Luciferase* gene as a normalization control (Fig. 4, C and D).

**Figure 4.**
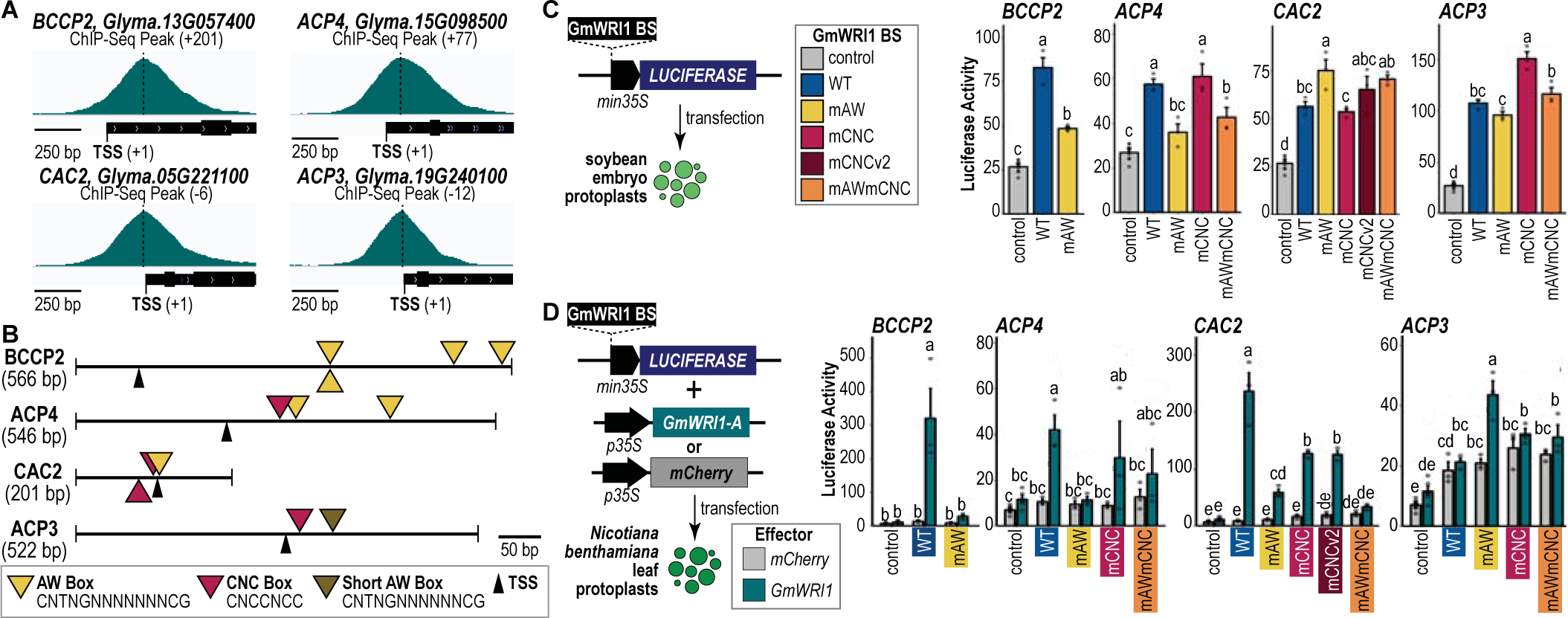
Functional analysis of GmWRI1 direct target gene binding sites. (A) Genome browser view showing the position of GmWRI1 ChIP-Seq peaks for *BCCP2* (Glyma.13G057400), *ACP4* (Glyma.15G098500), *CAC2* (Glyma.05G221100), and *ACP3* (Glyma.19G240100). ChIP-Seq peaks (dashed lines) and TSS positions (tick mark) are indicated. Peak distance from the TSS in bp is shown. (B) Diagrams of GmWRI1 binding sites for genes in (a) used in the transient assay experiments. Yellow, pink and brown symbols, respectively, indicate the positions of AW Box, CNC Box, and short AW Box motifs. (C) Effect of WT GmWRI1 binding sites and of binding sites with mutations in all detectable AW Box (mAW), CNC Box (mCNC) or both (mAWmCNC) DNA motifs on reporter gene activity in soybean cotyledon embryo protoplasts. (D) Transactivation analysis of WT and mutant GmWRI1 binding sites in tobacco leaf protoplasts transfected with *p35S:GmWRI1* (teal) or *p35S:mCHERRY* (grey). For (C) and (D), average firefly to Renilla luciferase activity ratios were multiplied by 100 and plotted with the respective standard errors. Lower-case letters indicate the significance of comparisons based on ANOVA and post hoc Tukey test results.

Constructs were assayed either by transfecting the plasmids into soybean embryo protoplasts or by co-transfecting the plasmid into *Nicotiana benthamiana* (tobacco) leaf protoplasts along with another plasmid with a *35S:GmWRI1* or *35S:mCHERRY* gene (Fig. 4, C and D).

We showed that all WT GmWRI1 binding sites promoted reporter gene activity in soybean embryo protoplasts, and all except *ACP3* resulted in reporter gene activation in leaf protoplasts when co-transfected with *35S:GmWRI1* (Fig. 4, C and D). These results suggest that *cis-*regulatory elements within these binding sites are sufficient to recruit GmWRI1 and, perhaps, other TFs to activate gene transcription. We assayed binding sites with mutated DNA motifs to determine if the AW Box or CNC Box acted as *cis-*regulatory elements and obtained variable results. As shown in Fig. 4, C and D, mutations in the AW Box of *ACP4* and *BCCP2* binding sites severely reduced reporter gene activity in both embryo and leaf protoplasts. By contrast, mutations in the CNC Boxes of these binding sites did not reduce reporter gene activity. These results suggest that GmWRI1 function depends primarily on AW Box elements in *ACP4* and *BCCP2* genes. Mutations in the AW Boxes and CNC Boxes of *CAC2* and *ACP3* binding sites did not compromise their enhancer activity in soybean embryos. However, mutations in either motif of *CAC2* binding sites compromised reporter gene activation in leaf protoplasts (Fig. 4D). Because there is positional overlap between one of the two CNC Boxes and the AW Box in the *CAC2* binding site (Fig. 4B), it was necessary to establish that the negative effect of the CNC Box mutations on enhancer activity did not result from alteration of the AW Box. We showed that mutation of only the upstream CNC Box (mCNCv2) was sufficient to cause a reduction in reporter gene activity in transactivation experiments with GmWRI1 in leaf protoplasts (Fig. 4D). Furthermore, *CAC2* binding sites with mutations in both CNC Boxes and in the AW Box (mAWmCNC) resulted in a greater reduction in reporter gene activity than those resulting from mutations in either motif separately (Fig. 4D). This suggests that the two motifs act additively or synergistically in transcriptional regulation.

The *ACP3* binding site did not display enhancer activity in leaf protoplasts, and AW Box and CNC Box mutations in this binding site did not compromise its enhancer activity in embryo protoplast (Fig. 4, C and D). These results open the possibility that GmWRI1 occupancy and function might require other TFs. We created a series of 100 bp overlapping deletions in the 500 bp region surrounding the *ACP3* ChIP-Seq peak and assayed the effects of deleting sequential 100 bp segments in embryo protoplasts to identify regions with *cis-*regulatory elements (SI Appendix Fig. S6, A and B). Although each deletion caused some reduction in reporter gene activity compared to the complete region, deletions of regions containing the GmWRI1 ChIP-Seq peak resulted in the greatest reduction in reporter gene activity (SI Appendix Fig. S6).

Because AW Box and CNC Box mutations in the *ACP3* binding site did not cause a reduction in reporter gene activity (Fig. 4, C and D) we conclude that other DNA elements are required for *ACP3* enhancer activity.

Together, these results suggest that the AW Box and CNC Box motifs in GmWRI1 binding sites function as enhancer elements. Although the majority of these *cis-*regulatory elements are located downstream of the TSS, they were able to promote GmWRI1 transcription when placed upstream of the TSS.

### GmWRI1 and GmLEC1 Control Fatty Acid-Related Genes through Distinct Binding Sites

In addition to FA and TAG biosynthesis genes, many GmWRI1 DT genes encode TFs (Dataset S1), several of which are involved in controlling major embryonic processes in soybean and/or Arabidopsis, such as LEC1, ABI3, FUS3, ABI4, AGL15, and WRI1 (32).

GmLEC1 is particularly interesting because it and AtLEC1 are major regulators of FA and TAG biosynthesis in embryos, and GmLEC1 binding sites have been profiled genome-wide (28, 29, 34). We previously showed that *GmWRI1-A* and *GmWRI1-B* are direct target genes of GmLEC1 in soybean embryos (28), and we show here that *GmLEC1-1* and *GmLEC1-2* are GmWRI1 DT genes (Dataset S1). These results suggest that GmWRI1 and GmLEC1 act in a transcriptional positive feedback circuit in the control of FA and TAG biosynthesis in embryos.

To understand how the two TFs regulate FA and TAG biosynthesis programs, we identified FAT GmWRI1 DT genes that are also bound by GmLEC1 in em-stage embryos (28). Fig. 5A shows that 69 of 111 of FAT GmWRI1 DT genes were also bound by GmLEC1 (p <1.5e-07). The GmWRI1 DT/GmLEC1 bound genes encode the vast majority of enzymes required for FA and TAG biosynthesis, although some key steps in the pathways are targeted uniquely by each TF (SI Appendix Fig. S7). Specifically, the KASIII enzyme (Fig. 1A) is encoded by a GmWRI1 DT gene that is not bound by GmLEC1 (SI Appendix Fig. S7, Dataset S3). By contrast, genes encoding fatty acyl-ACP thioesterases B (FATB) and glycerol-3- phosphate acyltransferase (GPAT) are direct targets of GmLEC1 but not GmWRI1 (28). These results suggest that although complementary, the two TFs serve some specific roles in FA and TAG biosynthesis in embryos. Moreover, FAT mRNAs encoded by GmWRI1 DT/GmLEC1 bound genes were at higher levels than those encoded by GmWRI1 DT genes alone (Fig. 5B). Our results suggest that GmWRI1 and GmLEC1 serve additive and/or synergistic functions in FAT gene regulation in embryos.

**Figure 5.**
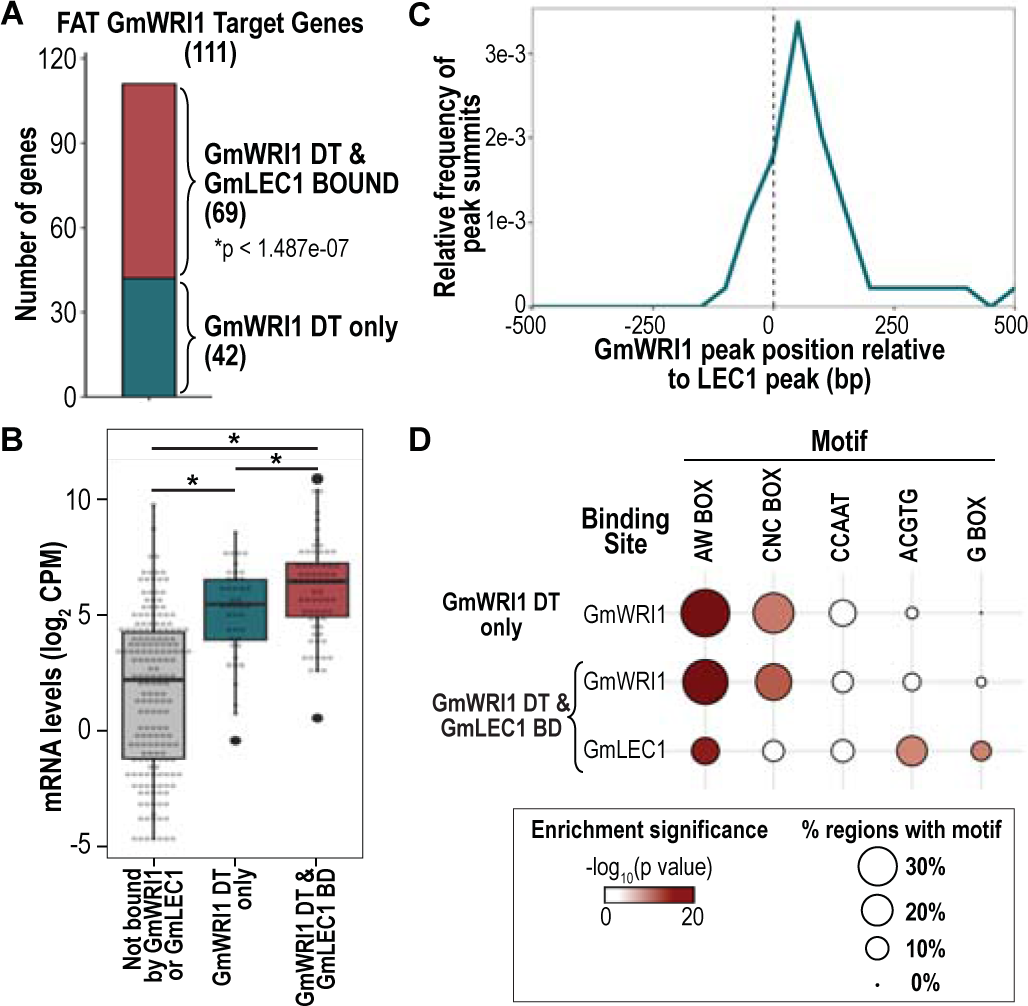
GmWRI1 and GmLEC1 act independently to control storage lipid biosynthesis in soybean embryos. (A) Bar graph shows the number of FAT GmWRI1 DT genes that are bound or not bound by GmLEC1. The *P* value significance of the overlap is indicated (hypergeometric distribution). (B) Box plots comparing the distribution of mRNA levels in em embryos (GEO accession GSE99571) of FAT genes are not bound by either GmWRI1 and GmLEC1 (grey), GmWRI1 DT genes that are not bound by GmLEC1 (teal), and GmWRI1 DT genes that are bound by GmLEC1 (dark red). Asterisk indicates significant differences between groups (Wilcoxon rank-sum test, p < 0.01). (C) GmWRI1 ChIP-Seq peak positions relative to LEC1 ChIP-Seq peaks (position 0) in FAT GmWRI1 DT genes that are also bound by GmLEC1. (D) DNA motif enrichment in GmWRI1 DT binding sites of genes that are not bound or bound by GmLEC1. Circle diameters depict the frequencies at which DNA motifs were identified in the indicated regions, and the color intensity indicates the statistical significance of the enrichment relative to the normal distribution of a population of randomly generated regions (Bonferroni- adjusted *P* values).

We next evaluated the positional relationship between GmWRI1 and GmLEC1 binding sites in FAT GmWRI1 DT genes. Fig. 5C shows that the median distance between GmWRI1 and GmLEC1 ChIP-Seq peak positions was 142 bp, ranging between 3 and 5,655 bp, suggesting that these TFs bind adjacent but distinct sites in target genes. To address this hypothesis, we analyzed DNA motif enrichment in a 50 bp window around ChIP-Seq peaks. As observed previously, GmWRI1 binding sites were strongly enriched for the presence of AW Box and CNC Box in the FAT genes targeted by GmWRI1 only and in the FAT GmWRI1 DT genes that are also bound by GmLEC1 (Fig. 5D). GmLEC1 binding sites were enriched for G Box-type DNA motifs (G Box and ACGTG) and the AW Box but not the CNC Box motifs (Fig. 5D). Although GmLEC1 has not been shown to bind G Box-type DNA motifs directly, its occupancy in FAT GmWRI1 DT genes is likely anchored by a bZIP type TF that binds G Box elements and physically interacts with GmLEC1, as suggested previously (28, 35). Given the distance between binding sites and the absence of evidence indicating that the two TFs interact physically, GmWRI1 and GmLEC1 appear to act independently to promote the transcription of FAT GmWRI1 DT genes.

## Discussion

### GmWRI1 is a Central Regulator of Storage Lipid Biosynthesis in Embryos

Fatty acid and TAG biosynthesis in embryos requires the activity of several enzymes located in distinct cellular compartments. Expression of genes encoding these enzymes must be highly coordinated to ensure the rapid and efficient incorporation and storage of carbon as FAs and TAGs during seed development (Fig. 1A)(3, 4). Given the critical role played by WRI1 in regulating FA biosynthesis genes and in controlling storage lipid accumulation in embryos (36), we obtained the first genome-wide profile of WRI1 binding sites and of the genes directly regulated by WRI1 in any plant.

Several lines of evidence indicate that GmWRI1 DT genes are directly regulated by GmWRI1 and that GmWRI1 is a key regulator of storage lipid biosynthesis. First, GmWRI1 DT genes correspond to FA and TAG related genes that are upregulated by ectopic GmWRI1 overexpression in soybean. Specifically, 24 of 55 genes upregulated by ectopic *GmWRI1* expression in transgenic soybean seeds, 19 of which are FAT genes, are GmWRI1 DT genes (25). Similarly, 7 of 18 genes, 6 of which are FAT genes, upregulated by ectopic *GmWRI1* in soybean roots are GmWRI1 DT genes (20). Second, there is significant overlap between GmWRI1 DT genes and homologous Arabidopsis genes that are downregulated in *atwri1* mutants (Fig. 2). We note that a microarray analysis of ∼3,500 Arabidopsis gene identified 25 downregulated genes in *atwri1* mutant when compared to WT seeds (37), 11 of which overlapped with the 1,213 list of downregulated genes in *atwri1* mutant embryos identified here (Dataset S5). This restricted overlap likely reflects the limited gene numbers assayed previously and reduced sensitivity of the analysis resulting from the use of RNA from whole seeds rather than embryos given that *AtWRI1* is primarily expressed embryonic tissues (38). Third, Arabidopsis gene homologs of GmWRI1 DT genes overlapped significantly with those that are upregulated upon ectopic overexpression of GmWRI1 in Arabidopsis leaf protoplasts (Fig. 2, Dataset S3). Fourth, GmWRI1 DT genes are enriched for FAT genes and encode the vast majority of enzymes involved in the FA and TAG biosynthetic pathways from the synthesis of malonyl-CoA in plastids to the assembly of TAG in the endoplasmic reticulum (Fig. 1, A and E).

Others have reported that AtWRI1 regulates genes encoding glycolytic enzymes (17, 19, 37). Hexoses funneled through the glycolysis pathway account for much of the carbon used for FA biosynthesis. Although glycolytic process was not an enriched GO term for GmWRI1 DT genes, seven genes encoding plastid-localized glycolytic enzymes were identified as GmWRI DT genes (Dataset S1). Together, these results indicate that GmWRI1 is a major regulator of storage lipid biosynthesis.

### Biological Relevance of GmWRI1 Binding Preference for Intragenic Regions

GmWRI1 binds preferentially with intragenic regions, consistent with the observation that AW Box and CNC Box motifs are enriched primarily downstream of the TSS (Figs. 3A, S1B, and S4). We demonstrated that intragenic binding was significant transcriptionally by showing that GmWRI1 binding sites with AW and CNC motifs located primarily downstream of the TSS in *BCCP2, ACP4*, and *CAC2* genes promote GmWRI1-mediated transcription when placed upstream of a minimal promoter (Fig. 4, C and D). Together, these results establish that the intragenic binding sites contain transcriptional enhancer elements.

Our finding that GmWRI1 DT genes possess intragenic binding sites is not unique. A survey of Arabidopsis TF binding sites revealed that 13 of 27 TFs analyzed exhibited substantial binding to intragenic sites (39), and an estimated 36% of enhancer elements in Drosophila genes are located in intragenic regions (40). A mechanistic explanation of how intragenic binding affects transcription comes from a recent study of the cucumber TENDRIL IDENTITY (CsTEN), a TCP TF of the CYC/TB1 clade. CsTEN binds primarily to target gene intragenic regions resulting in chromatin disassembly and transcription activation (41). CsTEN possesses an amino-terminal intrinsically disordered region domain with histone acetyltransferase activity that facilitates chromatin accessibility during target gene transcription. Interestingly, the AtWRI1 TF also has an amino-terminal intrinsically disordered region domain (42), opening the possibility that CsTEN and WRI1 share mechanistic similarities, although further experiments are needed to demonstrate such associations. Nevertheless, our results provided new insights into the mechanisms by which GmWRI1 functions in storage lipid biosynthesis in embryos.

### GmWRI1 Function is Dependent on AW Boxes and CNC Boxes

Our results provide compelling evidence that the AW Box serves as an enhancer element that mediates transcriptional regulation by WRI1. Mutations in AW Box elements in *BCCP2, ACP4,* and *CAC2* binding sites compromised their enhancer activity in transactivation assays with GmWRI1 in leaf protoplasts. Consistent with these findings, others have shown that AW Box elements are enriched in WRI1 target genes in several plants, that WRI1 binds AW Box elements *in vitro*, and that mutations in the AW Box compromised transcriptional activation by WRI1 (19, 20, 23, 25, 43–45).

A novel finding of our study is that DNA elements other than AW Boxes are involved in GmWRI1 transcriptional activation. Our finding that AW Box mutations did not compromise the enhancer activity of *CAC2* and *ACP3* binding sites in soybean embryo assays and of *ACP3* binding sites in leaf protoplast assays indicates that other DNA *cis* elements are present in these binding sites (Fig. 4, C and D). Consistent with this conclusion, only ∼55% of binding sites for FAT GmWRI1 DT genes contain full AW Boxes, implying that GmWRI1 may bind other DNA elements or that other TFs bind GmWRI1 binding sites.

Mutational analysis of the *CAC2* binding site in leaf cells demonstrated conclusively that the CNC Box in the GmWRI1 binding site is an enhancer element that promotes GmWRI1 transcriptional activity (Fig. 4D). The *CAC2* binding site also possesses AW Box enhancer elements that promote transcription. Comparison of the effects of AW Box and CNC Box mutations singly and in combination suggests that the two elements act additively or synergistically to enhance GmWRI1-mediated transcription. Consistent with the identification of the CNC Box as an enhancer element, the CNC Box is highly enriched in regions downstream of the TSS in GmWRI1 DT and AtWRI1 DR genes (Fig. S4 and S5). Enrichment of the CNC Box in GmWRI1 binding sites opens the possibility that the CNC Box may enhance GmWRI1 occupancy at these binding sites.

The mechanism by which the CNC Box enhances GmWRI1 transcriptional activity is not known, but similarities between the CNC Box and known DNA motifs provide potential clues about its function as an enhancer element. The CNC Box shares strong similarity with the CTCCGCC DNA motif that acts as an intragenic, positive transcriptional enhancer element that is bound by the CsTEN TCP-like TF (41). One model is that a TCP-like TF physically binds with the CNC Box and enhances GmWRI1 occupancy and functions in an AW Box-independent manner (Fig. 6). Recent studies showed that AtWRI1 interacts physically with the TCP TFs, AtTCP4, AtTCP10 and AtTCP24, although these TFs appear to repress seed oil accumulation (46) An alternate hypothesis is based on the fact that WRI1 is a member of the AP2/EREPBP TF family (16, 44). A survey of the Arabidopsis cistrome revealed that several AP2/EREBP domain TFs, such as ERF11, 15 and 105 and RAP212, bind DNA motifs similar to the CNC Box (47) It is possible that GmWRI1 could potentially bind directly with the CNC Box, although WRI1 has not been reported to bind physically with DNA motifs other than the AW Box. Additional experiments are needed to define the role of CNC Box elements in controlling the transcription of FAT genes regulated by GmWRI1 (Fig. 6).

**Figure 6.**
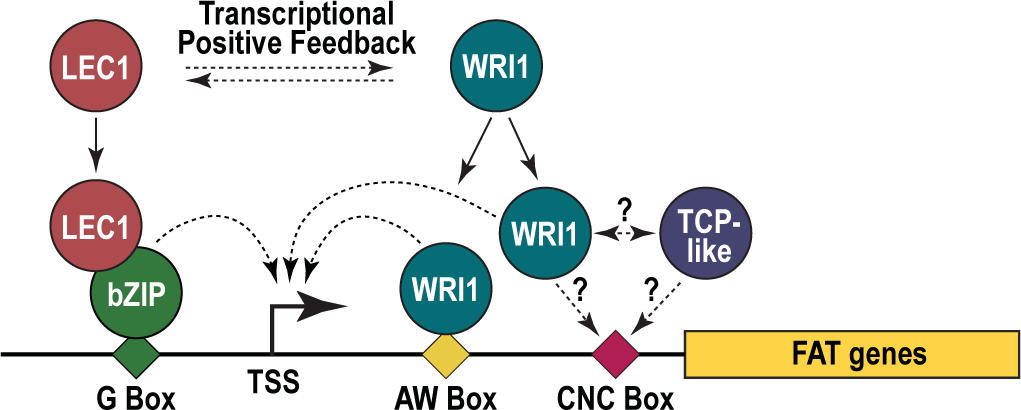
Model for the regulation of storage lipid accumulation in embryos. WRI1 and LEC1 act in a transcriptional positive feedback subcircuit to regulate FA and TAG biosynthesis in the embryo. GmWRI1 regulates FA and TAG-related gene expression by binding with AW Box and, possibly, CNC Box elements located downstream of the TSS. An alternative model is that another TF, such as a TCP-like TF, binds the CNC Box, potentially by interacting with WRI1. GmLEC1 binds FAT genes at sites that are distinct from GmWRI1 binding sites. GmLEC1 interacts with bZIP TFs that bind with G Box DNA elements located most frequently in regions upstream of the TSS. Dashed arrows indicate transcriptional activation

### GmWRI1 Acts with GmLEC1 and Other Transcription Factors to Establish a Regulatory Environment that Promotes Storage Lipid Accumulation in Embryos

In addition to establishing that GmWRI1 directly regulates genes encoding the vast majority of enzymes in the FA and TAG biosynthetic pathways, our results also indicate that GmWRI1 acts to alter the regulatory state of soybean embryos to promote storage lipid accumulation. We showed that GmWRI1 directly regulates several other well characterized TFs in seeds, including GmLEC1, GmFUS3, and GmABI3, whose counterparts in Arabidopsis transcriptionally regulate genes involved in storage lipid accumulation (34, 48). We also showed that GmWRI1 is a GmWRI1 DT, indicating that it is a self-activating TF. Thus, GmWRI1 is a critical component of a transcriptional regulatory network that promotes storage lipid accumulation and other processes during seed development.

The relationship between GmWRI1 and GmLEC1 provides a particularly illuminating example of the regulatory interplay that occurs between these TFs. Here, we showed that GmLEC1 is directly regulated by GmWRI1, that overexpression of GmWRI1 in Arabidopsis leaf protoplasts results in the upregulation of *AtLEC1*, and that *AtLEC1-LIKE* is downregulated in *atwri1* mutant embryos (Dataset S5). We showed previously that GmWRI1 is directly regulated by GmLEC1, that overexpression of *AtLEC1* results in elevated expression of *AtWRI1*, and that *AtWRI1* is downregulated in *atlec1* mutant seeds (28, 29). These findings indicate that WRI1 and LEC1 act in a positive feedback subcircuit to control FA and TAG biosynthesis in embryos through reciprocal regulation of the other TF’s expression and through their regulation of FA and TAG biosynthetic genes (Fig. 6). Positive feedback gene circuits act in gene regulatory networks to trigger irreversible changes in transcriptional programs (49). A hallmark of the onset of the seed maturation program is the *de novo* initiation of many biological programs that are silenced during the morphogenesis phase of the embryo development, such as storage lipid and protein accumulation (50). Initiation of the FA biosynthetic program in seeds requires a robust and stable regulatory state to ensure proper expression of genes required for the quick and efficient conversion of carbon into oil bodies. In this sense, a positive feedback subcircuit between GmLEC1 and GmWRI1 is likely to be required for the rapid onset of the FA and TAG accumulation programs during seed development.

GmWRI1 also appears to act in positive feedback circuits with GmABI3 and GmFUS3, which are both GmWRI1 DT genes (Dataset S1). GmABI3 directly regulates *GmWRI1*, and ectopic *GmABI3* expression activates *GmWRI1* (28, 51). Although GmFUS3 DT genes have not been identified, AtFUS3 directly regulates *AtWRI1* in Arabidopsis (52). These findings emphasize that GmWRI1 is an integral component of the transcriptional regulatory networks controlling storage lipid accumulation.

## Materials and Methods

### Chromatin Immunoprecipitation and DNA Sequencing

Soybean cv Williams 82 (W82) plants were grown, and embryos were isolated from seeds staged as described previously (29). GmWRI1 ChIP assays were conducted as described previously (29), with minor modifications. After nuclei purification, nuclei were resuspended in shearing buffer (10mM Tris pH8, 1mM EDTA, 0.1% SDS, 1mM PMSF), and chromatin was sheared to ∼250 bp fragments using a Covaris E220 ultrasonicator. After shearing, Triton X-100 and NaCl were added to chromatin to final concentrations of 1% and 150mM, respectively. Anti- GmWRI1 (Glyma.08G227700 and Glyma.15G221600) antibodies for ChIP were raised in rabbits against the peptide FSTQPGDHETESDVN, corresponding to the amino acids 352-366 of GLYMA.08G227700 and amino acids 348-361 (FSTQPGDETESDVN) of Glyma.15G221600.

Antibody reactivity was assessed using Western blot analysis (Dataset S2). ChIP DNA library preparation and DNA sequencing was done as described previously (29).

### RNA-Sequencing Experiments with Arabidopsis Leaf Protoplasts Overexpressing *GmWRI1* and with *atwri1* Mutant Embryos

For *GmWRI1* overexpression assays with Arabidopsis leaf protoplasts, three-week-old *Arabidopsis thaliana* Col-0 plants were grown as described (53). Arabidopsis leaf mesophyll protoplasts were isolated and transfected as described previously (28). For each biological replicate, 500,000 cells were transfected either with 10 µg of a plasmid carrying the *GmWRI1-A* cDNA that was synthesized using primers listed in Dataset S6, Table S1, and driven by the *CaMV35S* promoter (*p35S:GmWRI1*) or with a *35S:mCHERRY:NOSt* plasmid as a negative control. Protoplasts were incubated for 16 hours at room temperature following transfection.

Homozygous *atwri1-1* and WT Col-0 seeds were plated on GM media and grown on soil as described previously (53). Flowers from six-week-old plants were tagged at anthesis, and embryos were dissected from seeds 8 days after pollination (25-30 embryos per biological replicate, three biological replicates per genotype).

Total RNA was isolated using the RNAqueous™-Micro Total RNA Isolation Kit, following manufacturer’s instructions. RNA samples were then treated with DNAse for digestion of DNA using the TURBO DNA-free™ Kit, following manufacturer’s instructions. RNA-Seq libraries were prepared with Tecan Ovation RNA-Seq System V2 using 50 ng of total RNA following protocols as described by Kumar *et al.* (54), with NEXTflex ChIP-Seq Barcodes (BioScientific). DNA was sequenced with a NovaSeq 6000 sequencer.

### Cloning and Mutation of GmWRI1 Binding Sites

GmWRI1 binding sites (200 to 500 bp around the WRI1 ChIP-seq peak) were PCR amplified from the soybean genomic DNA and cloned as described previously (28). Mutated derivatives of binding sites and *ACP3* deletion series were synthesized by Twist Bioscience and inserted into pDLUC15 using the same cloning strategy. Mutant derivatives of *ACP4* were created by amplifying the WT *ACP4* binding site with primers containing the mutated sequences. Primers and the full DNA sequences of WRI1 binding sites and mutant derivatives are listed in Dataset S6, Table S1 and S2.

### Transient Expression Assays

Transient assays of firefly and Renilla luciferase activity in protoplasts were performed as described previously (28). All trials were done with three biological replicates, and one-way or two-way ANOVA with Tukey’s post hoc tests (P < 0.05) were used for the statistical analysis in soybean embryo protoplasts and tobacco leaf protoplasts, respectively.

### Data Analysis

#### GmWRI1 ChIP-Seq in cot and em stage soybean embryos and identification of GmWRI1 DT genes

ChIP-Seq data were analyzed as described previously (28), with the exception that MACS3 software (https://github.com/macs3-project/MACS) was used in place of MACS2.

Quality assessment of the ChIP-seq libraries were determined by evaluating the non-redundant fraction (NRF) and strand cross-correlation (CC), following ENCODE guidelines for ChIP-Seq data (26). NRF and CC values are summarized in Dataset S2 Because GmWRI1 binding sites were located primarily in regions downstream of the TSS, we designated genes bound by GmWRI1 as those with a reproducible ChIP-Seq peak within a 1 kb window upstream of its TSS or within the annotated gene body.

GmWRI1 DT genes are defined as those bound by GmWRI1 and coexpressed by GmWRI1 (Dataset S1). Because *GmWRI1* is expressed primarily in embryonic cells (Fig. 1C), we used the Harada-Goldberg Soybean Seed Development LCM RNA-Seq Dataset (GEO accessions, GSE57606, GSE46096, and GSE99109) to identify genes that are two-fold upregulated (FDR < 0.01) in embryo cotyledon abaxial or adaxial parenchyma versus seed coat hilum and seed coat parenchyma subregions. We then defined WRI1 co-expressed genes as those upregulated in at least two of the above comparisons (Dataset S1).

#### Identification of soybean FAT genes

To define FAT genes, we used the soybean Plant Metabolic Network database, SoyCyc, and extracted the soybean genes associated with the pathways listed in the Dataset S3. We also included genes annotated with the GO category for fatty acid biosynthetic process (GO:0006633).

#### RNA-Seq data from Arabidopsis protoplasts and embryos

Sequenced reads were demultiplexed and reads corresponding to rRNA sequences were removed. The resulting filtered reads were mapped to Arabidopsis primary transcripts (TAIR10) using bowtie v0.12.7 with parameters -v 2 -5 10 -3 4 -m 1 --best –strata. The EdgeR package (v3.10.5) was used to obtain normalized expression values using the Trimmed Mean of M-values (TMM) method, and to identify differentially expressed genes between the different genotypes (FDR < 0.05).

#### GO enrichment analysis

GO term enrichment analyses were done as described previously (28, 29). GO functional annotation was extracted from TAIR10 and Soybase databases and used for Arabidopsis and soybean datasets, respectively. A q value threshold of 0.01 was used to identify enriched GO categories.

#### DNA motif analysis

DNA motif *de novo* discovery analyses of binding sites in GmWRI1 DT genes and FAT GmWRI1 DT genes was performed using the MEME-ChIP tool from the MEME suite v5.3.3 as described previously (28), using a motif maximum length of 15 and a E-value cutoff of 0.01. Discovered motifs were compared to the Arabidopsis DAP-Seq TF and the top-ranked DNA motifs identified are listed in Dataset S4.

DNA motif enrichment analysis was performed as described (28), using the DNA motifs AW-BOX (CNTNGNNNNNNNCG), CNC-Box (CNCCNCC), G-Box (CACGTG), RY (CATGCA), and CCAAT-Box (CCAAT) to screen the GmWRI1 and GmLEC1 binding sites and 500 bps windows around the TSS of WRI1 directly regulated genes in Arabidopsis using the “known” function from HOMER (55).

## Accession Numbers

Data are available at GEO, under the accessions: GSE152310 and GSE253104 (GmWRI1 ChIP-seq at cot and em stages, respectively), GSE253109 (*atwri1* mutant embryo RNA Seq), and GSE253107 (GmWRI1 overexpression RNA Seq).

## Acknowledgments

We thank Makenzie Melby, Sana Mohammad, and Mary Irani for their technical assistance. This work was supported by a grant from the National Science Foundation Plant Genome Research Program to RBG and JJH.

